# Long-term exposure to polyamines leads to bacteriophage resistance in *Pseudomonas aeruginosa*

**DOI:** 10.64898/2026.07.14.738440

**Authors:** Raymond Finnerty, Christina Lim, Patrick R. Secor, Christopher W. Marshall

**Affiliations:** Department of Biological Sciences, Marquette University, Milwaukee, WI; Department of Microbiology and Cell Biology, Montana State University, Bozeman, MT

## Abstract

Bacteria often evolve resistance to phage infection by altering the cell-surface structures required for viral adsorption. However, the role extracellular metabolites play in influencing phage susceptibility and the evolution of phage resistance remains unclear. Here, we evaluated whether sustained exposure to putrescine, a polyamine released during phage-mediated cell lysis, alters susceptibility of the pathogen *Pseudomonas aeruginosa* to the type IV pili-dependent phage DMS3vir. Using adaptive laboratory evolution over ∼66 generations, we evolved *P. aeruginosa* with or without putrescine and with or without DMS3vir. As expected, direct phage exposure rapidly led to complete phage resistance. Interestingly, populations exposed to putrescine also developed phage resistance by the end of the experiment, despite having never encountered the phage. Whole-population genome sequencing revealed parallel mutations in genes associated with type IV pili and the global transcriptional regulator *mexT*. Using transposon insertion mutants in the type IV ATPases *pilT* and *pilB*, we confirmed that disruption of these genes leads to DMS3vir phage resistance. We also used a type IV pilus biogenesis factor *fimV* transposon mutant, which showed a putrescine-dependent reduction in phage susceptibility. These findings show that sustained exposure to a host-derived metabolite can drive the evolution of phage resistance through modification of key phage-adsorption sites and regulatory genes. Our work identifies elevated polyamine exposure as a selective pressure that promotes type IV pili-mediated phage resistance, even in the absence of phage exposure.

**IMPORTANCE:** *Pseudomonas aeruginosa* is a major cause of hospital-acquired infections and a key priority for phage-based therapies. Previous work has shown that the polyamine putrescine is released into the extracellular environment during cell lysis. These signals can then transiently reduce susceptibility to bacteriophage infection and alert neighboring cells to danger. Our research demonstrates that long-term exposure to putrescine can drive heritable phage resistance without prior exposure to phage. We show that resistance is linked to mutations in genes involved in type IV pili assembly. This work further demonstrates the critical role that polyamines can play in promoting phage resistance in bacterial communities.

## INTRODUCTION

Bacteriophages (phages) are viruses that infect bacteria and are the most abundant biological entity on the planet (1–3). Due to phages relying on bacterial hosts for replication, bacteria have evolved diverse mechanisms to prevent phage infection, replication, and spread (2, 4, 5). These resistance mechanisms are important for understanding bacterial evolution in both environmental and clinical settings, as well as for developing phage-based therapies. Phage therapeutics are being explored as a potential targeted treatment option that could work against a variety of antibiotic-resistant bacterial infections and improve patient outcomes (6). The rising interest in phage therapy is due to the increasing prevalence of antibiotic-resistant bacterial strains in many parts of the world (7, 8). A key advantage of phage therapy over broad spectrum antibiotics is the specificity of phages to their target bacteria, ensuring that only pathogenic bacteria are targeted with minimal collateral damage to the wider microbiota (9–11). A major barrier to phage therapy is that bacteria rapidly develop resistance to phages through multiple defense mechanisms (4, 5, 12). Therefore, understanding the factors that shape phage susceptibility and resistance is critical for both microbial ecology/evolution and the therapeutic use of phages.

*Pseudomonas aeruginosa* is one of the most concerning bacterial pathogens because its prevalence and antibiotic resistance are increasing in many parts of the world (13, 14), emphasizing the need to develop new therapies to treat antibiotic-resistant *P. aeruginosa* infections. *Pseudomonas aeruginosa* has a variety of phage defense systems and high phenotypic plasticity, allowing rapid adaptation to its environment (15–17). These factors make it a particularly challenging pathogen to deal with in clinical settings, emphasizing the need for novel treatment methods.

One recently described mechanism for phage resistance in *P. aeruginosa* is the release of intracellular molecules that can serve as an early warning system of phage lysis in neighboring cells (20). Specifically, this study showed that short-term exposure to the polyamine putrescine can inhibit DMS3vir phage replication (20). Polyamines are small, positively charged organic molecules with two or more amino groups that are found universally throughout nature (18, 19). Putrescine is the simplest polyamine and is found at a wide variety of concentrations in bacterial cells (0.1mM-30mM) (19). Polyamines have a variety of effects on bacterial physiology, including the promotion of biofilm production, elevating levels of c-di-GMP, and can influence phage susceptibility (19–21). Polyamines have also been shown to bind to and condense viral DNA through charge interactions (22, 23). These findings raise the possibility that elevated intracellular levels of polyamines could condense injected phage DNA, inhibit viral replication, and allow subpopulations of bacteria to evade phage lysis (20).

In this study, we investigated the effects of long-term putrescine exposure on *P. aeruginosa*. Our research investigates what genetic and phenotypic changes would be associated with prolonged putrescine exposure. Prior work has shown that short-term putrescine exposure transiently reduces phage susceptibility in *P. aeruginosa*, but whether sustained putrescine exposure leads to heritable phenotypic changes remained unclear (20). We wanted to expand on this work and use experimental evolution to understand the potential pathways involved in putrescine regulation and signaling in the cell. In this study, we demonstrated that exposure to elevated levels of putrescine leads to evolved, heritable phage resistance in *P. aeruginosa* without the population ever having been exposed to phage.

## RESULTS

### Long-term exposure to putrescine drives phage resistance

Extracellular polyamines can reduce phage susceptibility in *P. aeruginosa* (20), however the effects of sustained polyamine exposure on *P. aeruginosa* phage resistance are poorly defined. To examine the effects of long-term putrescine exposure on *P. aeruginosa*, we propagated *P. aeruginosa* strain MPAO1 every 24 hours for 10 days (∼66 generations) in each of four conditions: MPAO1 alone (control), MPAO1 with the virulent phage DMS3vir (VIR), MPAO1 with putrescine (PUT), and MPAO1 with putrescine and DMS3vir (PUT+VIR). Previous studies have shown that levels of intracellular putrescine ranged from 0.1-30 mM and that exogenous levels of putrescine can reach 50 mM following phage infection (20).

Therefore, 50 mM putrescine was added each day to mimic the extracellular conditions present when neighboring cells undergo lysis. Taken together, this experimental plan was designed to provide insight into the heritable genotypic and phenotypic effects that long-term exposure to elevated levels of putrescine would have on *P. aeruginosa*.

Each day, *P. aeruginosa* populations were exposed to a DMS3vir phage concentration of 10^8^ plaque forming units (PFU)/mL. This daily concentration of 10^8^ PFU/mL was empirically chosen to avoid *P. aeruginosa* population extinction while still allowing enough selective pressure to drive phage resistance. Previously, it was demonstrated that a DMS3 titer of 10^8^ PFU/mL was on the border between total population extinction and insufficient phage lysis (24).

Population sizes remained relatively stable throughout the entire experiment except for a slight drop at day 2 in the PUT+VIR condition (Fig 1A). This consistent population size indicates that neither 50 mM putrescine nor the daily addition of 10^8^ plaque-forming units (PFU/mL) of DMS3vir significantly reduced MPAO1 growth during the experiment. In contrast, DMS3vir phage titers in the VIR and PUT+VIR were ∼10^10^ PFU/mL after the first 24 hours but were reduced to 10^8^ PFU/mL and 10^7^ PFU/mL by day 10, respectively (Fig 1B). This decline in phage titer suggests that the entire population of *P. aeruginosa* evolved resistance to DMS3vir, thereby limiting phage replication over the course of the experiment.

**Fig 1.**
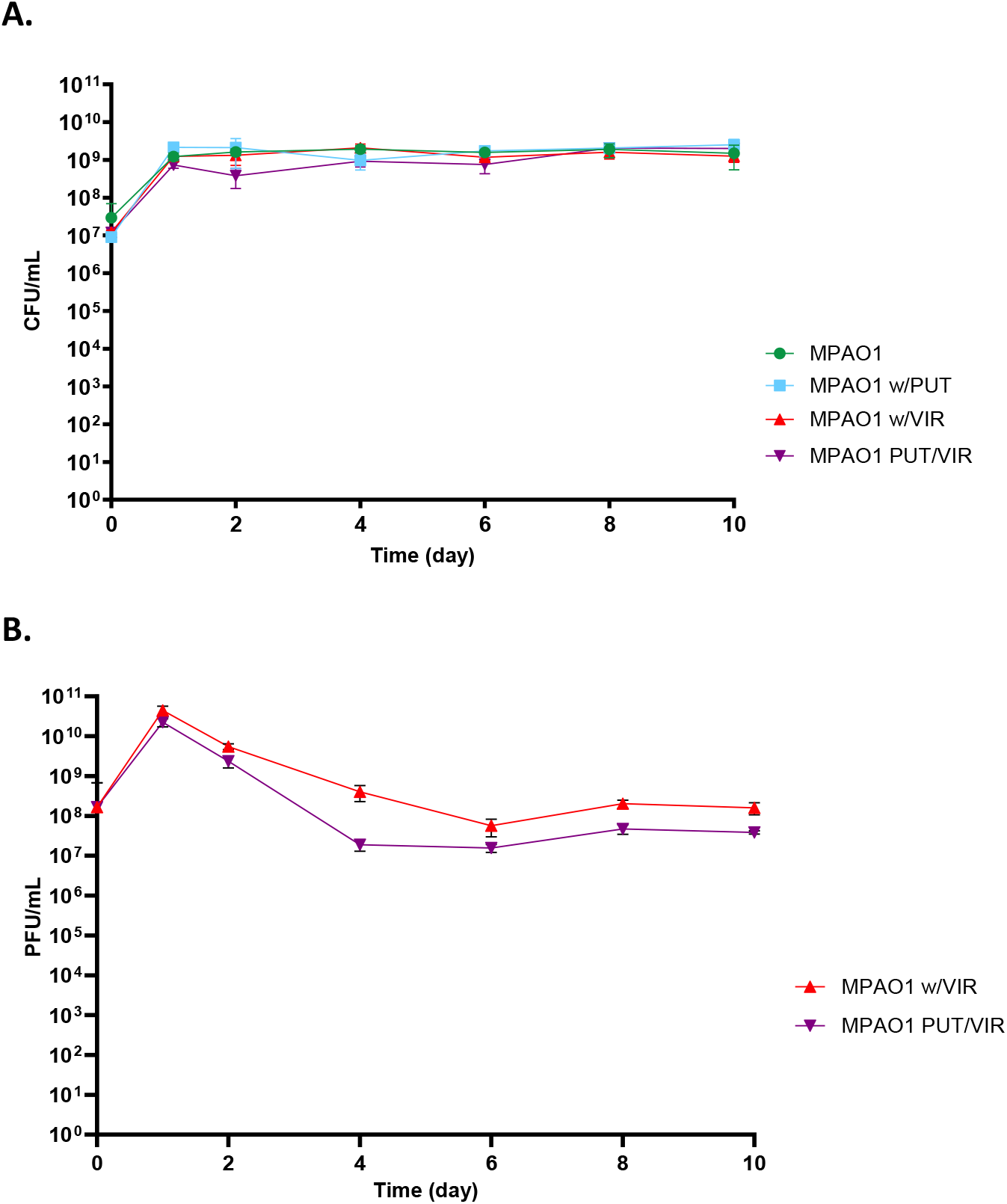
Population sizes of *P. aeruginosa* (A) and phage titers of DMS3vir (B) over the 10-day experiment.

We next tested to see if the evolved lineages could be infected by DMS3vir on day 3 or day 10. All control MPAO1-only evolved lineages were susceptible to phage infection, as indicated by the ∼10^10^ PFU/mL phage titers across all timepoints (Fig 2). All populations exposed to phage (PUT+VIR and VIR) were resistant to phage infection (below detection limit) by day 3 and remained so for the duration of the experiment (Fig 2). Surprisingly, the PUT evolved lineage acquired resistance by day 10 of the experiment (Fig 2). This indicates that prolonged exposure to the polyamine putrescine is sufficient for phage resistance even in the absence of phage exposure.

**Fig 2.**
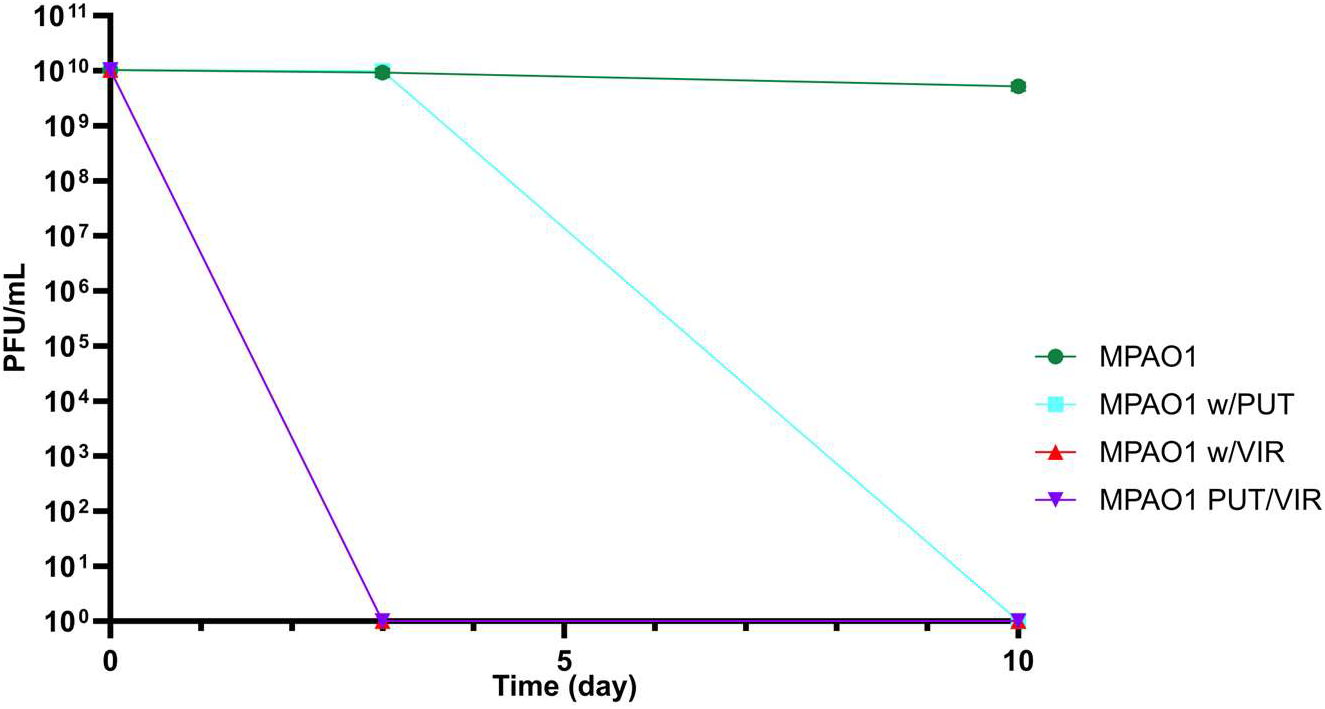
Phage susceptibility of evolved populations at days 0, 3, and 10 of the evolution experiment. Tested in the absence of putrescine.

### Mutations in type IV pili genes mediate phage resistance

To identify the mutations that led to the observed phage resistance in the PUT only exposed populations, we did whole-population genome sequencing on samples from days 3 and 10 of the experiment. While we observed sporadic, low frequency mutations (<10% of the total population) in a few *pil* genes in PUT at day 3, we observed several high-frequency mutations in genes involving type IV pili (T4P) in PUT-only populations on day 10 of the experiment (Table 1, Supplemental Table 1). T4P are filamentous appendages that are observed in many bacteria; these appendages support twitching motility, biofilm formation, and are a common target for phage recognition and adsorption (25, 26).

**Table 1.**
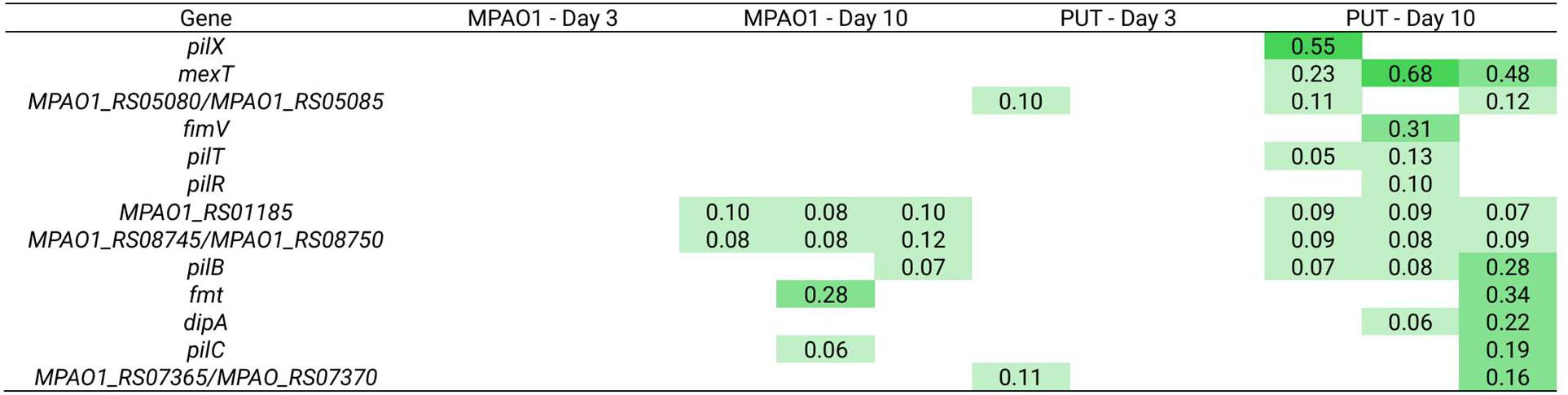
Genes containing mutations at indicated frequencies. Mutations are listed if the sum of mutations in that gene were greater than or equal to 5% of the population. Frequencies listed are the sum of all mutations in that gene in each population. Mutations in three replicate populations per condition on days 3 and 10 are shown.

Mutations in the T4P conferring phage resistance are in line with the literature that states that DMS3vir uses T4P for adsorption of *P. aeruginosa* (26, 27). Consistent with this model, if PUT exposure confers phage resistance, this resistance likely derives from the site of phage adsorption, the T4P. Therefore, the observation of high-frequency mutations in T4P at day 10 and not day 3 is consistent with our data showing PUT populations were resistant to phage at day 10 but not day 3 (Fig 2).

We also observed that the PUT populations had parallel mutations in the transcriptional activator *mexT* (Table 1). MexT positively regulates the efflux pump MexEF-OprN and a large number of other genes throughout the genome (28, 29). However, T4P have not been found to be differentially regulated by MexT knockout mutants (30) and so we wanted to test whether mutations in *mexT* could reduce phage susceptibility as is implied by our mutational parallelism in the PUT treatment coupled with the observed phage resistance by day 10 (Fig 2). We used transposon mutants with insertions in the *mexT* gene in addition to transposon mutants of *fimV, pilB*, and *pilT* and incubated the isogenic mutants with DMS3vir (Fig 3A). When enumerating PFUs, we found that the *mexT* transposon mutant was not significantly less susceptible to DMS3vir in the presence of putrescine compared to the ancestor (Fig 3B), indicating that mexT mutations might be adaptive during putrescine exposure independent of phage resistance mechanisms. However, the transposon insertion mutants of *fimV, pilB*, and *pilT* were all resistant to DMS3vir in putrescine. These mutants are expected to be representative of the putative loss of function mutations observed in the evolved lineages (Table 1). Based on these results, phage resistance can be explained by the combinations of mutations in the T4P in all the PUT-evolved populations.

**Fig 3.**
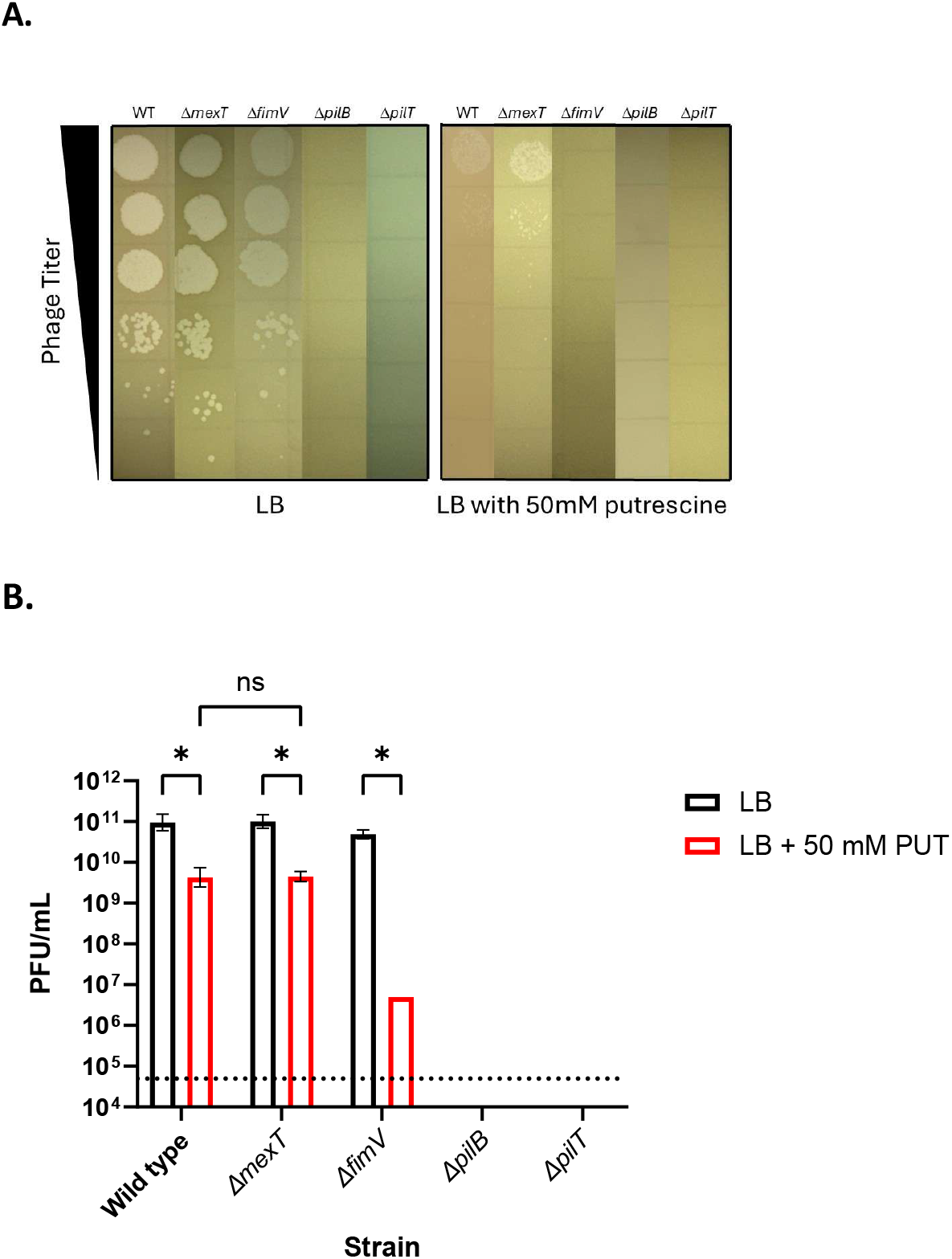
Phage plaques for the ancestor MPAO1 (WT) and four different transposon insertion mutants in the presence and absence of 50 mM putrescine. Phage titers are 10^6^ PFU/mL - 10^1^ PFU/mL of DMS3vir from top to bottom (A). PFU/mL of DMS3vir on LB agar or LB agar supplemented with 50 mM putrescine with lawns of specified strains. Faint clearance zones were observed at 5×10^6^ PFU/mL for Δ*fimV*. No clearance zones were observed below the limit of detection (5×10^4^ PFU/mL) for Δ*fimV*, Δ*pilB*, and Δ*pilT*. Results are the mean ± SD of three biological and technical replicates, ANOVA *P<0.05.

## DISCUSSION

Previously, it was demonstrated that supplementation of exogenous putrescine reduces phage susceptibility in *P. aeruginosa*, a finding we also observe (Fig 3A, B) (20). Uniquely, we have shown that after nearly 66 generations of exposure to high levels of exogenous putrescine, populations of *P. aeruginosa* evolved heritable genetic phage resistance to the phage DMS3vir (Fig 2). This long-term exposure to putrescine selected for mutations in genes associated with T4P, the primary site of DMS3vir adsorption (Table 1) (26, 27). Therefore, the *P. aeruginosa* response to putrescine is complex, where short-term exposure acts as an early warning system of phage infection that can reduce susceptibility to phage (20) and where long-term exposure to putrescine in the absence of phage can lead to phage resistance even when putrescine is removed (Fig 2).

We observed mutations in T4P genes across PUT-exposed populations (Table 1). This is consistent with previous work demonstrating that resistance to T4P-dependent phages can be mediated through inactivating mutants in several T4P genes (31). One of the putrescine-exposed evolutionary replicates had a mutation in the minor pilin gene *pilX*. PilX is thought to provide structure to the tip of the T4P (32– 34). Due to the structure provided by minor pilins, it is likely that loss of function may lead to decreased adsorption ability of DMS3vir. In a separate replicate population, we observed a mutation in *pilR*. PilR is a key transcriptional regulator that modulates levels of the major pilin PilA in *P. aeruginosa* (35, 36).

Previous studies have demonstrated that reduction of *pilA* in other bacterial species leads to reductions in biofilm formation, motility, virulence, and phage susceptibility (37, 38). Based on this, we expect that mutations in *pilR* could lead to a reduction in expression of *pilA* and therefore contribute to phage resistance (31).

In two independent PUT-evolved populations, we observed *pilB* and *pilT* mutations. PilB functions as an ATPase for the T4P (39-42), and loss of function mutations in *pilB* are associated with impaired T4P assembly (40). Therefore, we suspect that our putrescine exposed evolved lineages with mutations in *pilB* also lack the ability to assemble T4P, becoming resistant to phage infection. Further supporting this, transposon insertion mutants of *pilB* were completely resistant to the DMS3vir phage (Fig 3). PilT is a retraction ATPase that depolymerizes T4P (40, 43, 44). PilT loss of function mutants are still able to assemble T4P, but the retraction ability of Δ*pilT* mutants is inhibited (25, 40). It has been previously reported that retraction of pili is a likely mode for infection by T4P-dependent phage (45). Based on these previous studies and our work here, the combination of mutations observed in T4P are sufficient to explain why putrescine-exposed populations evolved resistance to the DMS3vir phage.

Interestingly, a mutation in *fimV* occurred in one of the PUT-only exposed populations. FimV is a key regulatory protein that has been shown to interact with many T4P proteins across multiple domains such as the alignment proteins PilM/N/O/P and secretin PilQ (46, 47). Inactivating mutants of *fimV* can lead to a nonmotile phenotype (47). However, these nonmotile mutants were still susceptible to infection by pilus-dependent phages (47). In contrast, a transposon insertion mutant of *fimV* provided resistance to phage in the presence of putrescine (Fig 3). This finding explains how this population could be phage resistant, though the exact mechanism of this resistance is unknown.

Despite the strong evidence of adaptation that mutational parallelism in evolving populations provides (48), it is still unclear why the mutations in T4P were adaptive in the absence of phage. It has been shown that putrescine can promote elevated cyclic-di-GMP levels in *P. aeruginosa* (21). Cyclic-di-GMP is a key regulator in the switching of *P. aeruginosa* from a planktonic to biofilm lifestyle (49). Cyclic-di-GMP has been shown to enhance bacterial binding through elevating production of exopolysaccharides (EPS) that allow bacteria to adhere to surfaces (50, 51). One aspect of high biofilm formation is that the oxygen levels through the depth of the biofilm and in the surrounding environment can rapidly deplete (52, 53) and it has been shown that T4P can no longer retract under conditions of high cyclic di-GMP and no oxygen (51). Therefore, altering the T4P structure and function under these conditions may be adaptive. It is also well-established that T4P is regulated by another secondary messenger, cyclic-AMP (55), but there appears to be scenarios where dynamic levels of cyclic-di-GMP and cyclic-AMP can regulate biofilm and T4P depending upon external stimuli (55–57). Based on this, we propose that there is a relationship between elevated levels of putrescine indirectly selecting mutations in T4P, possibly through a mechanism involving cyclic-di-GMP levels. This could be particularly emphasized in one of the PUT replicate populations containing a mutation in the phosphodiesterase gene *dipA* (Table 1) where deletion mutants are deficient in attachment and have elevated cyclic-di-GMP levels .

Mutations in *mexT* appear in all replicates treated with PUT. It has been previously reported that an overexpression of *mexT* leads to a down regulation of PA2592, a hypothetical polyamine binding protein (29). It is possible that *mexT* mutations occur in populations exposed to putrescine to maintain higher levels of PA2592 to better respond to elevated polyamine levels, potentially conferring a fitness advantage. Although we admit that this hypothesis is speculative. Another possible reason for *mexT* mutants in this study is its possible regulatory role in nitrosative stress (58). High concentrations of polyamines may lead to stress conditions in the cell that could be mitigated by regulatory networks under the influence of *mexT* (29, 30). More work will need to be done to understand why *mexT* mutations are adaptive in putrescine and how that might contribute to reduced phage susceptibility.

In this study, we have shown that long-term exposure to the polyamine putrescine is sufficient to develop resistance to the T4P-dependent bacteriophage DMS3vir. Phage resistance is likely achieved through a combination of mutations in genes implicated in the development and function of the T4P. Future studies elucidating the mechanism by which putrescine exposure selects for mutations would provide critical insight into how bacterial communities resist phage infection and respond to danger signals. Studies into the role of *mexT* in high putrescine environments or other nitrogen-rich environments would also provide valuable information on this broad transcriptional activator. Finally, understanding whether putrescine or other polyamines are sufficient to achieve phage resistance in other species of bacteria could help develop models for general phage defense mechanisms.

## MATERIALS AND METHODS

### Bacterial and Phage Strains, Growth Conditions

*Pseudomonas aeruginosa* MPAO1 (ASM1610748v1) and the bacteriophage DMS3vir (24) were used for adaptive laboratory evolution experiments. Lysogeny broth (LB) (Lennox; Fisher Bioreagents, catalog no. BP1427500) and agar were used to revive cultures from freezer stocks, propagate, and conduct plaque assays. A 1 M stock solution of putrescine dihydrochloride (MP Biomedicals) was filter sterilized using a 0.22 µm filter. This 1M solution was added to molten LB agar (1.5% agar) or LB broth and mixed to achieve a final concentration of 50 mM putrescine for all experiments. To test the possible effects of mutations in certain genes, we used transposon-insertion mutants from the Manoil transposon mutant library of MPAO1 (59). The strains used were PW1223 (mexT-E04::ISlacZ/hah), PW6238 (fimV-F01::ISphoA/hah), PW8623 (pilB-G07::ISlacZ/hah), and PW1728 (pilT-E12::ISphoA/hah).

### Phage Propagation

High titer (∼10^9^ PFU/mL) DMS3vir was generated by infecting a host culture of PAO1 that was grown in 6 mL trypticase soy broth for 6 hours. 300 µL of DMS3vir stock was incubated with host culture overnight. Aliquots of propagated DMS3vir were added to microcentrifuge tubes and spun down for 10 min at 20000 x g. Supernatant was passed through a 0.22 µm filter to remove bacterial debris. Phage supernatant was diluted 10-fold in Phosphate Buffer Saline (PBS). 10 µL of appropriate phage dilution (10^-5^ to 10^-8^), 5 mL LB soft agar (0.6% agar), and 200 µL of fresh 6 h PAO1 host culture were combined and poured over LB agar plate (100 mm x 15 mm, 1.5% agar). The soft agar top layer was allowed to solidify, and then plates were incubated at 37°C overnight. PFUs were counted to determine the titer of the new phage stock.

### Adaptive Laboratory Evolution

Freezer stocks of *P. aeruginosa* MPAO1 were thawed and streak plated on LB agar and incubated overnight at 37°C. A single colony was resuspended in 5mL LB, grown at 37°C overnight, and the overnight culture was used to inoculate 20 tubes containing 5mL of LB. The tubes were divided into 5 replicates each of 4 conditions: MPAO1 only (PAO1), MPAO1+50mM putrescine (PUT), MPAO1+10^8^ PFU/mL DMS3vir added every day (VIR), and MPAO1+50mM putrescine+10^8^ PFU/mL DMS3vir added every day (PUT+VIR). A 1:100 transfer (50µL into 5mL) of culture from the previous day was transferred every 24 hours +/- 1 hour into a new 5mL tube of LB. This continued for 10 days, which is equivalent to approximately 66 generations.

### CFU Sampling

On days 0, 1, 3, 4, 6, 8, and 10, viable cells were determined by colony-forming unit (CFU) assays. Cultures were mixed thoroughly, and samples were serially diluted 10-fold in PBS to final dilutions ranging from 10^-5^ to 10^-8^. 100 µL aliquots of each dilution were plated onto LB agar plates (100 mm x 15 mm, 1.5% agar) and incubated overnight at 37°C. Colonies were counted to calculate CFU/mL.

### PFU Sampling

On days 0, 1, 3, 4, 6, 8, and 10, phage titers were determined by a plaque-forming unit (PFU) assay. 2 mL aliquots of all conditions were added to microcentrifuge tubes and spun down for 10 min at 20000 x g. Supernatant was passed through a 0.22 µm filter to remove bacterial debris. Phage supernatant was diluted 10-fold in PBS. 10 µL of appropriate phage dilutions (10^-3^ to 10^-8^), 5 mL LB soft agar (0.6% agar), and 200 µL of fresh 6 h PAO1 host culture were combined and poured over LB agar plate (100 mm x 15 mm, 1.5% agar). The soft agar top layer was allowed to solidify, and then plates were incubated at 37°C overnight. PFUs were counted to determine the titer of the phage present throughout evolution.

### DNA extraction and sequencing

DNA was extracted from the evolved populations and the ancestor using the DNeasy Blood and Tissue kit according to the manufacturer’s protocol (Qiagen). Purified DNA was sent to SeqCoast Genomics for library preparation and sequencing. Samples were prepared for sequencing using the Illumina DNA Prep tagmentation kit. Samples were sequenced to a minimum of 1 Gbp per sample on an Illumina NextSeq2000 using 2x150bp reads. Sequence reads were trimmed and filtered using trimmomatic v.0.39 (60) with Leading, Trailing, and sliding window quality scores all set to 20. Trimmed forward and reverse paired reads were used to call variants against the MPAO1 reference (GCF_016107485.1) using breseq v.0.39 with the –p flag specified (61, 62).

### Phage Plaquing Assay

Phage plaquing assays were conducted following an established protocol with slight modifications (63). Bacteria were grown overnight at 37°C in LB media with and without 50 mM putrescine. Overnight cultures were subcultured 1:100 in either LB or LB with 50 mM putrescine and grown at 37°C for 3 hours. After 3 hours, the subculture was normalized to an OD_600_ of 0.3. Then 320 µL of normalized culture was combined with 8 mL of LB soft agar (0.6% agar) with or without 50 mM putrescine. Soft agar was poured over square LB agar plates with or without 50 mM putrescine. These plates were air dried in a biosafety hood for 15 minutes with lids off. DMS3vir phage stock was serially diluted in LB 1:10 to achieve a concentration range of 10^8^-10^1^ PFU/mL. 5 µL of each phage concentration was spotted onto the solidified soft agar. Plates were once again allowed to air dry in a biosafety cabinet with the lids off for 15 minutes. They were then inverted and incubated at 37°C overnight. Plaquing assays were repeated three times.

## Data Availability

Sequence reads were deposited to the NCBI SRA under BioProject PRJNA1458946 with accession numbers SAMN57530720 - SAMN57530732.

## ACKNOWLEDGEMENTS

We thank the Department of the Army, US Army Engineer Research and Development Center award number W9132T-24-2-0007. We also thank the Pseudomonas aeruginosa transposon mutant library funded by NIH grant #SINGH19R0.

## SUPPLEMENTAL MATERIAL

Supplemental Table 1. Detailed information about all mutations observed throughout the experiment.

